# Transcriptome-Wide Off-Target Effects of Steric-Blocking Oligonucleotides

**DOI:** 10.1101/2020.09.03.281667

**Authors:** Erle M. Holgersen, Shreshth Gandhi, Yongchao Zhou, Jinkuk Kim, Brandon Vaz, Jovanka Bogojeski, Magdalena Bugno, Zvi Shalev, Kahlin Cheung-Ong, João Gonçalves, Matthew O’Hara, Mark Sun, Boyko Kakaradov, Andrew Delong, Daniele Merico, Amit G. Deshwar

## Abstract

Steric-blocking oligonucleotides (SBOs) are short, single-stranded nucleic acids designed to modulate gene expression by binding to mRNA and blocking access from cellular machinery such as splicing factors. SBOs have the potential to bind to near-complementary sites in the transcriptome, causing off-target effects. In this study, we used RNA-seq to evaluate the off-target differential splicing events of 81 SBOs and differential expression events of 46 SBOs. Our results suggest that differential splicing events are predominantly hybridization-driven, while differential expression events are more common and driven by other mechanisms. We further evaluated the performance of *in silico* screens for off-target events, and found an edit distance cutoff of three to result in a sensitivity of 14% and false discovery rate of 99%. A machine learning model incorporating splicing predictions substantially improved the ability to prioritize low edit distance hits, increasing sensitivity from 4% to 26% at a fixed FDR. Despite these large improvements in performance, the approach does not detect the majority of events at a false discovery rate below 99%. Our results suggest that *in silico* methods are currently of limited use for predicting the off-target effects of SBOs.

## Introduction

Antisense oligonucleotides (ASOs) are short, single stranded nucleic acids designed to bind to a target mRNA using Watson-Crick base pairing (Khvorova & Watts, 2017; Roberts et al., 2020; X. Shen & Corey, 2018). They have been successfully used as therapeutics by downregulating gene expression (Benson et al., 2018) and altering splicing (Finkel et al., 2017).

As with other therapeutic compounds, ASOs can cause off-target effects. These can be grouped into effects caused by unintended hybridization to RNA regions that are similar to the ASO target sequence (known as hybridization-dependent off-target events), sequence-dependent effects resulting from ASO-protein interactions (W. Shen et al., 2018), and sequence-independent effects resulting from the chemical properties of the ASO or delivery system. Hybridization-driven effects depend on the sequence, and so it has been suggested that they can be identified through *in silico* screens for near-complementary sites in the transcriptome (Lindow et al., 2012).

Previous studies have assessed the hybridization-dependent off-target effects of ASOs designed to degrade the target mRNA through RNase H cleavage. Oligonucleotides acting through this mechanism are known as gapmers, due to the presence of DNA bases flanked by modified bases (Crooke, 2004; X. Shen & Corey, 2018). One study used microarrays to assess the off-target gene expression changes of two locked nucleic acid gapmers (Yoshida et al., 2019). They calculated the edit distance, a count of the number of mismatches or gaps between the ASO and RNA sequence, to all genes, and found that 139 / 256 (54%) of all downregulated genes were at an edit distance of zero or one. Others have used qPCR dose-response experiments to investigate off-targets nominated from *in silico* screens of 96 high viability gapmers (Watt et al., 2020). In this study 97/ 832 predicted off-target sites had a reduction in gene expression with potency within 10-fold of the intended target transcript.

Steric-blocking oligonucleotides (SBOs) are fully modified ASOs that do not contain any DNA bases. Without DNA bases, RNAse H does not recognize the binding between the oligonucleotide and mRNA, and so the target mRNA is not degraded. SBOs instead act by blocking access to regulatory proteins and modifying RNA secondary structure (Havens & Hastings, 2016; Rigo et al., 2012). These properties have been exploited therapeutically to modulate splicing (Dulla et al., 2018; Finkel et al., 2017; J. Kim et al., 2019; Komaki et al., 2018) or to increase expression (Lim et al., 2020).

Since not all potential binding sites overlap a regulatory element, hybridization-dependent off-target effects are expected to be less frequent than for gapmers. One group used reverse-transcriptase PCR to quantify splicing changes at off-target binding sites with low predicted minimum free energy, a measure of the stability of the ASO/ RNA complex, and observed a splicing change for 22/108 exons (Scharner et al., 2020).

In this study, we use RNA-seq to comprehensively characterize the off-target splicing effects of 81 steric-blocking oligonucleotides. Differential expression off-target effects are characterized for a subset of 46 SBOs with sufficient biological replicates (see Methods). By assessing off-target effects transcriptome-wide, not only at near-complementary sites, we are able to evaluate the sensitivity and specificity of *in silico* methods. To our knowledge this is the most extensive characterization of off-target transcriptome changes induced by SBOs.

## Methods

### SBO design and synthesis

81 PS-MOE steric-blocking oligos of lengths 16 to 20 were synthesized at Integrated DNA Technologies using solid-supported methods in an oligonucleotide synthesizer. 2’-O-methoxyethyl (MOE) nucleotide phosphoramidites of adenosine (A), guanosine (G), 5-methyl cytidine (5-methyl C), and thymidine (T) were used in iterative detritylation, coupling, capping, and sulfurization steps. Protecting groups were removed from the oligonucleotide and the support was cleaved before desalting. The amount of salt-free oligonucleotide in the aqueous solution was measured using UV absorbance of the solution. The tube was then dried by vacuum and the resulting oligonucleotide pellet was resuspended in 1xTE buffer, pH 8.0, to a final concentration of 100 uM, using the known amount of salt-free oligonucleotide in the tube.

### RNA-seq experiments

105 RNA-seq experiments were performed in HepG2 cells, HEK293T cells, or PXB-cells, human hepatocytes isolated from a repopulated mouse liver (Hata et al., 2020). Each experiment compared samples transfected with a steric blocking oligo (SBO) to mock-transfected samples. Two to six (median three) biological replicates were used per condition. SBO concentrations were chosen to maximize the on-target effect for several therapeutic programs (data not shown).

Oligonucleotide transfections were performed by reverse transfection in HepG2 and HEK293T cells using Lipofectamine RNAiMAX reagent (ThermoFisher Scientific). Briefly, 300 pmol of oligonucleotide was mixed with 200 ul of OptiMEM reduced serum medium (ThermoFisher scientific) in a 12 well plate. 3 ul of RNAiMAX was then added to each well, mixed by rocking, and allowed to incubate for 20 minutes prior to addition of cells. Following the incubation, 3 × 10^5 cells in 800 ul of media without antibiotics (DMEM + 10% FBS for HepG2, IMDM + 10% Cosmic calf serum for HEK293T) was added to each well and mixed by rocking, giving a final oligonucleotide concentration of 300nM Cells were then incubated for 48 hours, after which RNA extractions were performed either manually using the Qiagen RNeasy Mini Kit (Qiagen) or by using the QIAcube automated extraction system according to the manufacturer’s recommended protocols (inclusive of DNase digestion steps).

For PXB-cells, oligonucleotide transfections were performed by forward transfection using Lipofectamine RNAiMAX reagent. Briefly, 6.5 × 10^4 PXB cells were first seeded in 96 well plates in a volume of 100 ul antibiotic free PXB culture medium plus 10% FBS. After incubation for 24 hours, 50 pmol of oligonucleotide was mixed with 20 ul of OptiMEM reduced serum media combined with 0.3 ul of Lipofectamine RNAiMAX and allowed to incubate for 20 minutes. Following Lipofectamine incubation, OptiMEM/Lipofectamine/oligonucleotide mixtures were added to the wells (417 nM final concentration of oligonucleotide), mixed by rocking, and returned to the cell culture incubator for 48 hours. For RNA extractions, the Qiagen RNeasy Mini Kit was used according to the manufacturer’s recommended protocol with the following exceptions. Initially, 100 ul of buffer RLT plus beta-mercaptoethanol was added to each well containing cells in the plate and pipetted up and down to lyse cells. 6 replicate wells with 100 ul of lysate were then combined together and an equal volume of 70% ethanol was added to the lysate. Two separate centrifugations were performed to process the entire volume of lysate/ethanol mixture. The remainder of the protocol was followed according to the manufacturer’s recommended conditions (inclusive of DNase digestion steps).

RNA quality was assessed using a Bioanalyzer 2100, and samples with RNA integrity score below 7.5 were dropped. RNA-seq libraries were prepared with the NEBNext Ultra II Directional RNA preparation kit, with polyA-selection performed using the NEBNext Poly(A) mRNA Magnetic Isolation module. Sequencing was done on an Illumina HiSeq 2500 or NovaSeq 6000 sequencer.

### Cell viability assay

HepG2 cells were reverse transfected with SBOs using RNAiMAX in 96 well plates. 20,000 HEPG2 cells were seeded per well. SBOs were transfected at 400 nM and cells were incubated for 48 hrs at 37°C and 0.5% CO_2_. Viability was assessed using the CellTiter-Fluor Cell Viability Assay (Promega) and performed in quadruplicate. To calculate SBO viability scores, background fluorescence was first subtracted from the fluorescence in all wells with SBOs. Negative and positive controls on each plate were then used to create a linear mapping to a reference dataset, ensuring that the reported values are comparable across experiments.

### RNA-seq analysis

Reads were aligned with HISAT2 v2.1.0 (D. Kim et al., 2019) to obtain full alignments for differential splicing analyses. The alignment index was generated by combining Gencode v25 annotations with Intropolis (Nellore et al., 2016) splicing junctions (filtered to junctions supported by at least two samples and five reads, with one end annotated in Gencode v25, and spliced in at least 0.01% of the time). The first 20 million reads for each sample were first aligned to detect novel splice sites, and these splice sites were used as input for a final HISAT2 run to align all reads. The samples had a median of 60.2 million mapped paired-end reads. Quality control was performed by assessing read coverage, percentage duplicated reads, and 5’ to 3’ read bias with FastQC v0.11.8 and RSeQC v2.6.4.

### Quantifying exon usage

We developed a novel method for quantifying exon usage based on spliced reads. For each exon, a set of possible upstream donors and downstream acceptors is compiled from all overlapping annotated Gencode v27 transcripts. Splicing junctions mapping from any of these splice sites to the exon boundaries are counted as inclusion reads, *I*, while splicing junctions mapping directly from upstream splice sites to downstream ones without including the exon are counted as exclusion reads, *E*. The splicing junctions themselves do not need to be consistent with an annotated transcript, but both the 5’ and the 3’ end of the junction must be annotated as a splice site.

Because every mRNA molecule can result in at most two inclusion reads and one exclusion read, the inclusion reads are divided by two. Percent spliced in (PSI) of the exon can then be calculated as the ratio of inclusion reads to total reads.

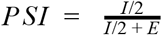

In the case where the exon of interest shares exactly one splice site with another exon, reads mapping to the end with the shared splicing junction cannot be used to distinguish between the two exons and are not counted. Only the end with the alternative splice site end is used, and spliced reads mapping to the alternative boundary of the other annotated exon are counted as exclusion reads. Since only one end of the exon is considered, each mRNA molecule can give rise to at most one inclusion read, and the inclusion read count is not divided by two.

If the exon shares its acceptor site with another exon, and its donor site with a different one, PSI cannot be quantified and such exons are dropped from the analysis. This was the case for 19,072 / 241,061 (7.9%) of unique exon intervals in Gencode v27.

### Detecting splicing changes

To test for significant differences in PSI between treated and control samples, we use a bootstrap test. If the observed difference in PSI is due to chance, it should be comparable to the differences we would see if the treated and control sample labels are shuffled. To simulate this scenario, we used a two-step bootstrap procedure that starts by randomly selecting samples with replacement, and then samples reads with replacement.

At each bootstrap iteration, treated and control samples are first sampled with replacement. The reads for these samples are pooled to a common set of reads. For each “treated” and “control” sample, reads are bootstrapped from this pooled set. The observed difference in PSI (dPSI) between the actual treated and control samples was compared to the simulated differences to obtain an empirical p-value. Each exon was initially run for 1000 bootstrap iterations. If the p-value was below 0.05, 49,000 additional bootstrap iterations were performed to obtain a more accurate p-value estimate.

### Detecting expression changes

To estimate transcript abundances for differential expression analysis, reads were pseudo-aligned to Gencode v27lift37 with Kallisto v0.46.0 (Bray et al., 2016), using 1000 bootstrap iterations and sequence-based bias correction. Transcript-level counts were aggregated to gene-level counts with tximport v1.10.1. DESeq2 v1.22.0 (Love et al., 2014) was used to test for differential expression between treated and control samples. Multiple testing correction was performed using the Benjamini-Hochberg method (Benjamini & Hochberg, 1995).

### Experiment reproducibility

Average transcripts per million and PSI-estimates were compared between repeated experiments using Spearman correlation. To assess reproducibility of differential expression and splicing results, log fold changes and dPSI were compared. We used the observed absolute effect size (log fold change or dPSI) as a predictor of significant events in the replication experiment. Sensitivity and false discovery rate was evaluated at different thresholds.

### Quantifying effects by edit distance

We computed the minimal edit distance between each SBO and different regions in the transcriptome. Edit distance was computed up to a maximum of four for SBOs of length 16 or 17 and a maximum of five for SBOs of length 18 or longer. Sites without any binding site at the maximum edit distance or lower were labelled as background.

To quantify splicing events by edit distance, we calculated the edit distance to all non-terminal in-frame exons, as skipping an out-of-frame exon is likely to trigger nonsense-mediated decay. We further removed all exons with a total read coverage less than 15 for either cases or controls due to low power (34,004-47,427, median of 36,889 filtered in-frame exons). Multiple testing correction was performed on the remaining exons using the Benjamini-Hochberg method. For each edit distance bin, we counted the proportion of exons where a significant (q < 0.05) change in splicing was observed.

To quantify differential expression events, comparisons with fewer than three case or control samples were removed due to low power, resulting in 64 comparisons of 46 unique SBOs. 32 of the SBOs were designed to hybridize to a specific region of the transcriptome (3’ UTR N = 5, exonic N = 20, intronic N = 7), five were designed as non-targeting controls, three as “promiscuous” SBOs with exact complementarity to many regions, and six to bind to RNA-binding protein motifs. We calculated the minimal edit distance between the SBO and the full gene (pre-mRNA or mature principal APPRIS v27 transcript), 5’ UTR, 3’ UTR, and out-of-frame exons. For each category, the proportion of significant changes in gene expression was calculated within each edit distance bin.

### Sequence-dependence of expression events

RNA-seq experiments were done with five SBOs targeting the same exon such that four SBOs overlap each other in two sets of two. Treated samples were compared to mock-transfected samples, and the Spearman correlation of log fold changes was computed for all pairs of SBOs. To compare the overlap of differential expression events with absolute log fold change greater than 1, the Jaccard index was computed for all pairs of SBOs.

### Predicting splicing changes

We evaluated the sensitivity and false discovery rate of different edit distance-based and energy-based methods for predicting off-target events. In-frame exons with a statistically significant change in PSI and an absolute dPSI of at least 0.5 were labelled as positive off-target hits. The delta G of SBO/ exon hybridization was computed using the RNA-cofold and RNA-plex methods included in ViennaRNA v2.4.14 (Lorenz et al., 2011), with a padding of 50 bp around the exon sequence. Exons longer than 1000 bp were excluded.

### Combined splicing effect and binding affinity model

To investigate the predictive ability of a model that factors in both splicing as well as binding affinity we trained a Gradient-Boosted Decision Tree (GBDT) using the LightGBM algorithm (Ke et al., 2017). We first created high quality splicing junction annotations using Gencode v27 and Intropolis (Nellore et al., 2016). For all genomic locations within protein-coding genes we labelled whether it lied within an intron or exon, or if it corresponded to an acceptor or donor splicing junction. We trained a Deep Convolutional Neural Network (CNN) that took the raw genomic sequence in a large window (16 kbp) and predicted the splicing annotations for every position in the input. We then used this CNN to predict the change in splicing caused by the SBO, assuming perfect hybridization to the exon. These predicted splicing scores combined with the predicted delta G, PSI of control samples, and features describing the relative position of the SBO with respect to the exon were then fed into the GBDT to predict the likelihood of an exon-SBO pair being an off-target hit. The model was trained and evaluated on exon-SBO pairs at an edit distance of five or lower (exon body or 200bp into flanking intron). We split the dataset with a 70:15:15 split for training,validation and testing ensuring that no SBO or exon appears in multiple splits. Having data splits fully disjointed by both SBO and exon sequence, we can get an unbiased estimate of the model’s generalization performance on new unseen examples.

## Results

We systematically assessed the off-target effects of splice-switching steric blocking oligonucleotides (SBOs) through 105 RNA-seq experiments with 81 different SBOs (**Supplementary Table 1**). 39 of the SBOs were designed to hybridize to specific places in the transcriptome (3’ UTR N = 5, exonic N = 25, intronic N = 9), a total of 18 different genes. Of the remaining 42, five were designed as non-targeting controls, 31 as “promiscuous” SBOs with complementarity to many parts of the transcriptome (2-30,670, median of 4 exact matches in the transcriptome), and six to be complementary to RNA-binding protein motifs (**Supplementary Figure 1**).

For each RNA-seq experiment, treated samples were compared to mock-transfected samples and putative off-target events were identified through differential splicing analyses (**Figure 1**, **Methods**). Differential expression events were assessed for a subset of 46 SBOs with sufficient number of replicates. The experiments had a median of 322.5 (range 3-5,403) differentially expressed genes (q-value < 0.05) with a reduction in expression of at least 50% or increase of at least 2x. Differential splicing events were less frequent, with a median of 5 (range 0-140) differentially used exons with a large change in PSI (|dPSI| > 0.5). As expected based on a hybridization and sequence-specific mechanism, non-targeting control oligos had the lowest numbers of differential splicing (median = 1) and differential expression (median = 96) events (**Table 1**).

**Figure 1:**
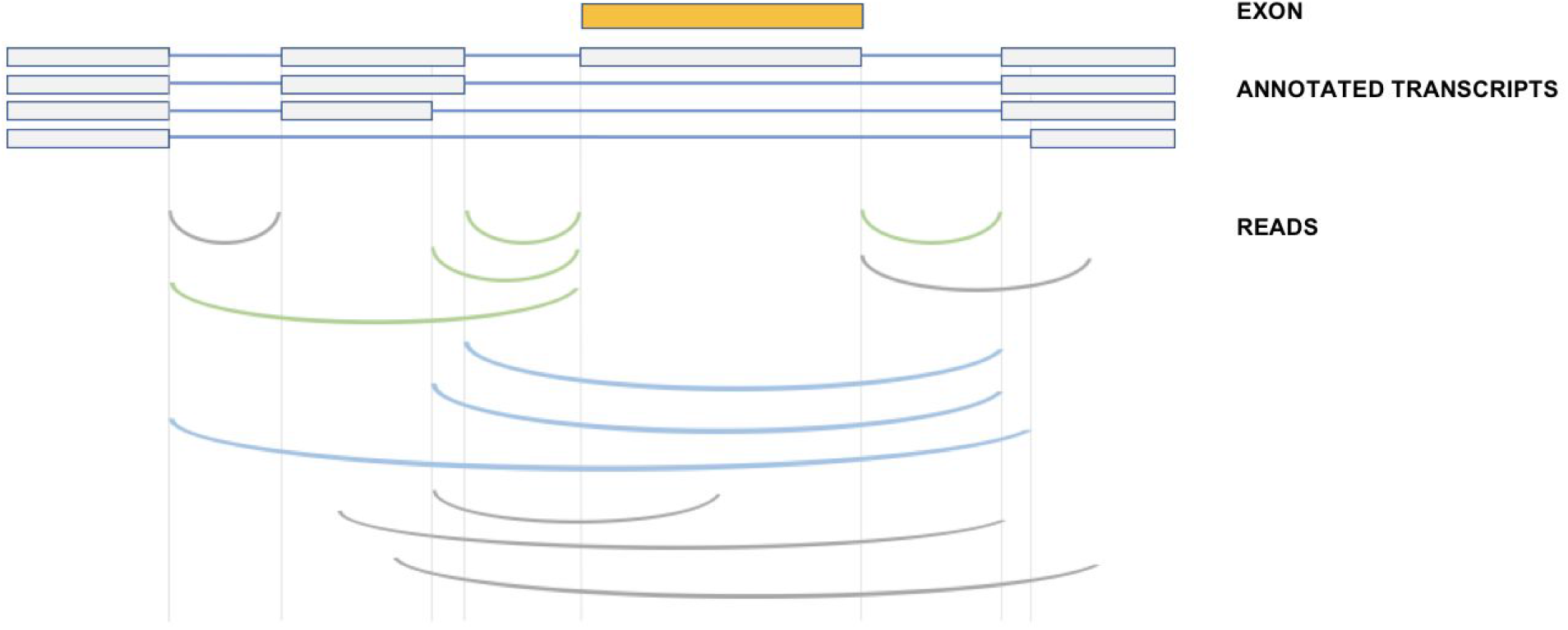
Quantification of percent spliced in for each exon. Overlapping transcripts are used to identify a set of upstream and downstream splice sites, and spliced reads mapping between these and the exon boundaries are counted as inclusion reads (green). Reads mapping directly between the upstream and downstream splice sites are counted as exclusion reads (blue). Reads where one or both ends does not match an annotated splicing junction are not counted (grey).

**Table 1:** Number of significant differential expression events (absolute log2 fold change > 1) and differential splicing events (absolute dPSI > 0.5) by category of SBO.

| Analysis | Category | Q1 | Median | Q3 | Number of experiments | Median replicates |
| --- | --- | --- | --- | --- | --- | --- |
| expression | 3' UTR | 639 | 857 | 1393 | 5 | 3 |
| expression | exonic | 63.75 | 397.5 | 1190.25 | 30 | 3 |
| expression | intronic | 29 | 137 | 750 | 9 | 3 |
| expression | non-targeting control | 7.75 | 95.5 | 456 | 8 | 3 |
| expression | promiscuous | 127 | 838 | 2263 | 6 | 3 |
| expression | RBP motif | 8.5 | 124.5 | 1244.75 | 6 | 3 |
| splicing | 3' UTR | 6 | 9 | 11 | 5 | 3 |
| splicing | exonic | 1.5 | 8 | 17 | 35 | 3 |
| splicing | intronic | 0 | 1 | 2 | 13 | 3 |
| splicing | non-targeting control | 0 | 1 | 2 | 9 | 3 |
| splicing | promiscuous | 2 | 6 | 21 | 37 | 2 |
| splicing | RBP motif | 0.5 | 2.5 | 31.5 | 6 | 3 |

We assessed the reproducibility of RNA-seq by conducting repeated experiments in HepG2 cells with four SBOs, each with four to six replicates. Transfections were performed on different days by different people, and sequencing done in separate batches. Average expression and PSI-estimates were highly reproducible (Spearman correlations of 0.88 to 0.91, **Supplementary Figure 2**), while differential results comparing cases to controls were more variable. 750 / 4788 (15.7%) differential expression events with a reduction of 50% or increase of 2x and 11 / 60 (18.3%) differential splicing events with a change in PSI of at least 0.5 were shared between the two experiments. If we consider all significant changes regardless of effect size, the overlap in differential expression events increases to 11,804 / 32,172. When using the observed changes in the first experiment to predict the significant events in the replication experiment, differential splicing is more predictable than differential expression (**Supplementary Figure 3**). In particular, differential splicing events with a large decrease in PSI are generally consistent between experiments. Several of these events occur at high edit distances (**Supplementary Figure 4**).

Next, we sought to quantify off-target events at near-complementary binding sites. To quantify the similarity between the SBO and off-target binding site, we counted the number of mismatches or gaps (edit distance) between the SBO and the reverse complement of the binding site. We computed the edit distance between the 81 SBOs and all in-frame non-terminal exons, and evaluated the probability of observing an off-target effect for edit distance zero to five. Exons without a low edit distance binding site were included in the analysis to provide an estimate of the background rate of events.

Splicing changes are strongly enriched at low edit distances, with decreasing probability of observing a difference and smaller effect sizes as the edit distance increases (**Figure 2a**). At exons that contain a binding site perfectly complementary to the SBO, the probability of seeing a change in PSI of at least 0.2 is around 35%, although this could be partially driven by the selection of SBOs with an on-target effect. This result is consistent with previous screens of steric-blocking oligos (Hua et al., 2007; Sinha et al., 2018), where not all exonic binding SBOs alter splicing.

**Figure 2:**
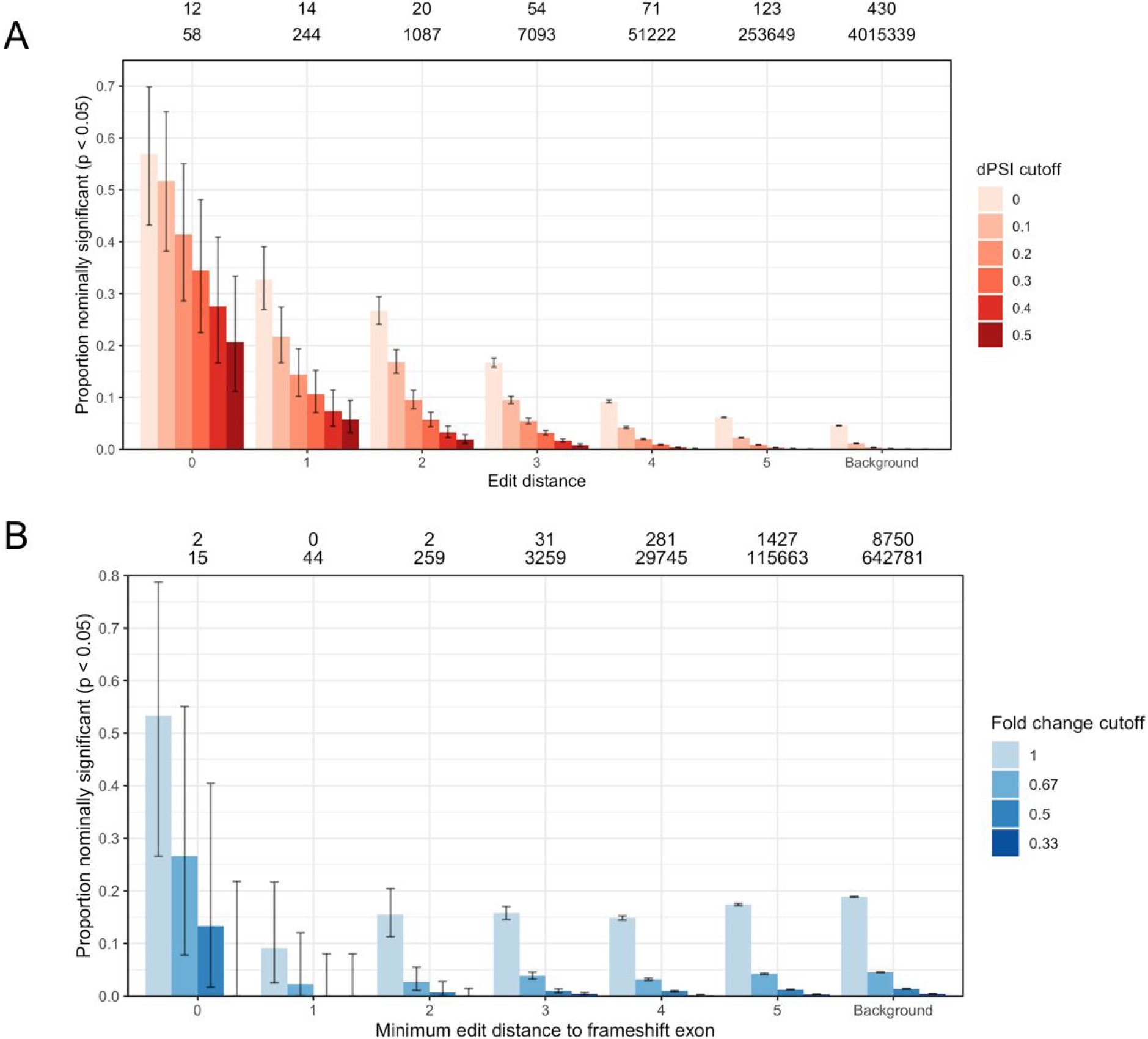
Proportion of potential binding sites resulting in a splicing (A) and expression (B) change, broken down by edit distance and effect size. The numbers above the bars give the number of significant (q-value < 0.05) events with absolute dPSI > 0.5 (A) or fold change < 0.5 (B) and the total number of events by edit distance. Error bars show 95% binomial proportion confidence intervals.

While the probability of observing a change in splicing drops off with more gaps and mismatches between the SBO and off-target binding site, even high edit distances have an enrichment of events compared to exons without an off-target binding site. Exons at edit distance five have a 5-fold enrichment of large splicing changes compared to background exons. Due to the larger number of candidate exons, most differential splicing events occur at high edit distances (**Supplementary Table 2**). 678 / 724 (93.6%) large splicing changes occur at exons with an edit distance of three or higher. Similar results are observed when restricting the analysis to the subset of 46 SBOs used for differential expression (**Supplementary Figure 5**).

By contrast, expression changes do not have a clear association with edit distance (**Figure 2b, Supplementary Table 3**). The background set of genes without a low edit distance hit have around a 19% probability of seeing a downregulation in expression, far higher than the expected type I error rate of 5%. Similar results were observed when looking at off-target binding sites in other regions of the gene and restricting to high viability SBOs (**Supplementary Figure 6–7**).

To assess whether differential expression events that do not depend on direct hybridization still depend on sequence, we used five SBOs designed to skip the same exon. The two pairs of overlapping SBOs had more similar expression changes than SBOs that do not overlap, although the concordance in differentially expressed genes is low for all pairs (**Supplementary Figure 8**). Based on these results, we chose to focus on differential splicing events as a more direct readout of hybridization-dependent effects.

For gapmer oligonucleotides, it has previously been reported that intronic binding sites are more susceptible to off-target effects than exonic regions (Kamola et al., 2015). To investigate the effects of intronic off-target binding for steric-blocking oligonucleotides, we repeated the differential splicing analysis when including the flanking intron (**Figure 3**). The enrichment of significant events at low edit distances drops off as intronic sequences are included, consistent with there being fewer splicing enhancer and silencer elements in intronic regions.

**Figure 3:**
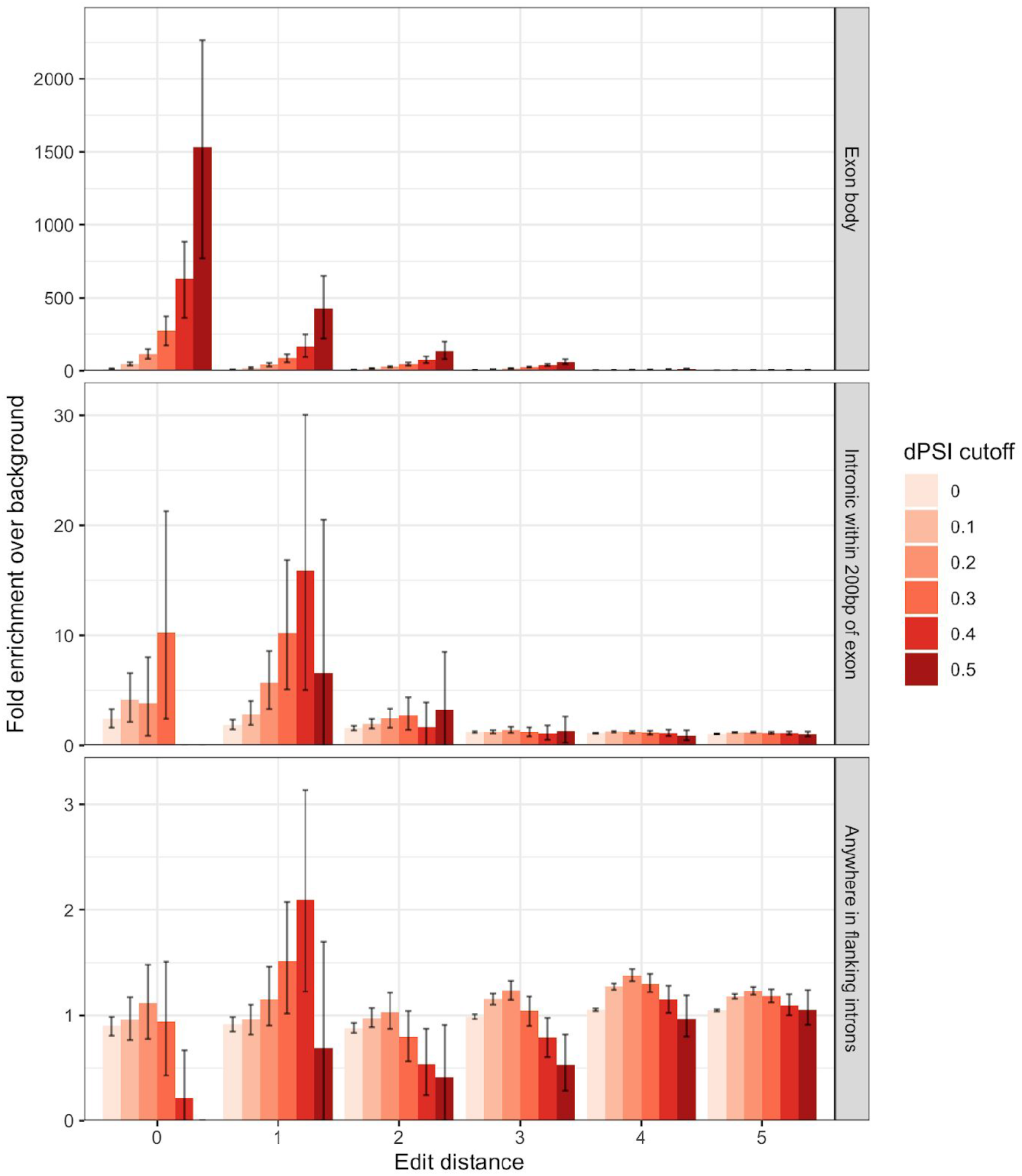
Enrichment of hits at different compared to a background of hits at edit distance 6+ at different dPSI cutoffs, broken by region. Error bars show 95% bootstrap confidence intervals.

The US Food and Drug Administration (FDA) draft guidance for Hepatitis B virus drugs suggested using *in silico* screens to identify potential off-target binding sites with three or fewer mismatches to an ASO (Food and Drug Administration Center for Drug Evaluation and Research, 2018). While splicing changes are more likely to occur if the edit distance between the SBO and exon is low, the majority of differential splicing events in our dataset still occur at higher edit distance sites. We therefore set out to evaluate the performance of different edit distance cutoffs for predicting off-target effects (**Figure 4**).

**Figure 4:**
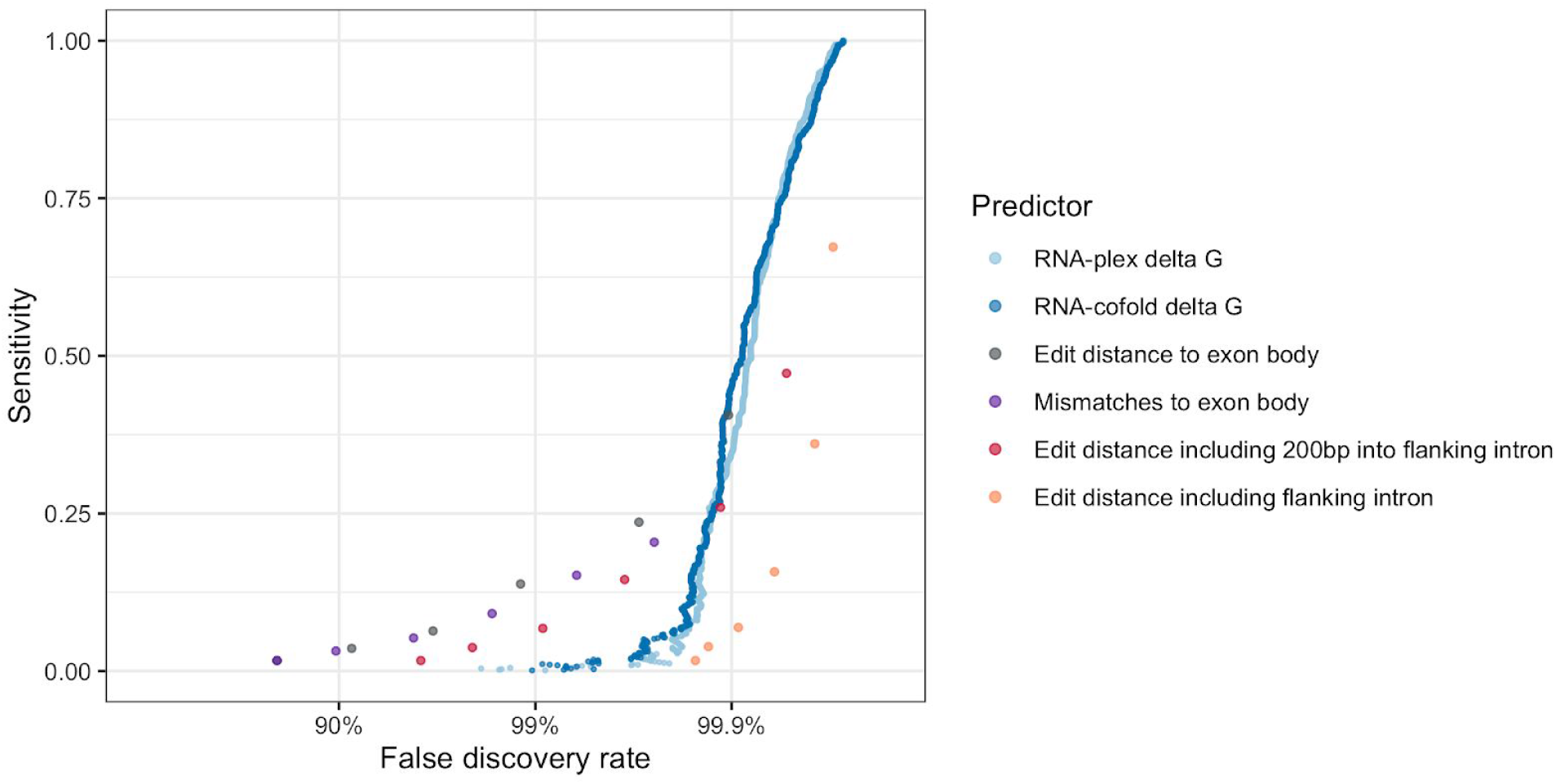
Performance of different predictors at identifying off-target splicing events with a change in PSI of at least 0.5.

If only exonic sites with exact complementarity are considered as potential off-target binding sites, only 1.6% of all large off-target splicing (absolute dPSI > 0.5) events are recovered at a false discovery rate of 80%. Increasing the edit distance cutoff improves sensitivity, but results in an overwhelming false discovery rate. Even at an edit distance of 5, only 40.6% of all differential splicing events are recovered, and 99.91% of predicted off-target sites do not have a change in splicing. We compared this performance to gapless edit distance, edit distance with flanking intronic sequence, and predicted minimum free energy (delta G) from RNA-cofold and RNA-plex.

None of the predictors are able to identify the majority of off-target events at a false discovery rate below 99%. Following the FDA guidance of considering off-target binding sites at three or fewer mismatches would only identify 9.1% of the true differential splicing events, and 98.3% of the identified sites would be false positives. Similar results are observed when removing the SBOs designed to hybridize to more than one region in the transcriptome (**Supplementary Figure 9**).

Edit distance and minimum free energy predictions measure the likelihood of an SBO binding to a particular sequence, but do not capture any information on the expected splicing effect. To explore whether *in silico* hits (edit distance five or lower) can be further prioritized based on the expected splicing effect, we trained a gradient-boosted tree (**Methods**) to predict splicing changes with a change in PSI of 0.2 or more, using both splicing predictions and binding affinity predictions as input. This model shows a clear improvement over binding affinity alone when evaluated on a test set of unseen SBOs and exons, increasing the sensitivity at 90% false discovery rate from 4.4% to 26.1% (**Figure 5**). Due to the lack of a clear association with edit distance, we did not attempt to train a similar model for differential expression results.

**Figure 5:**
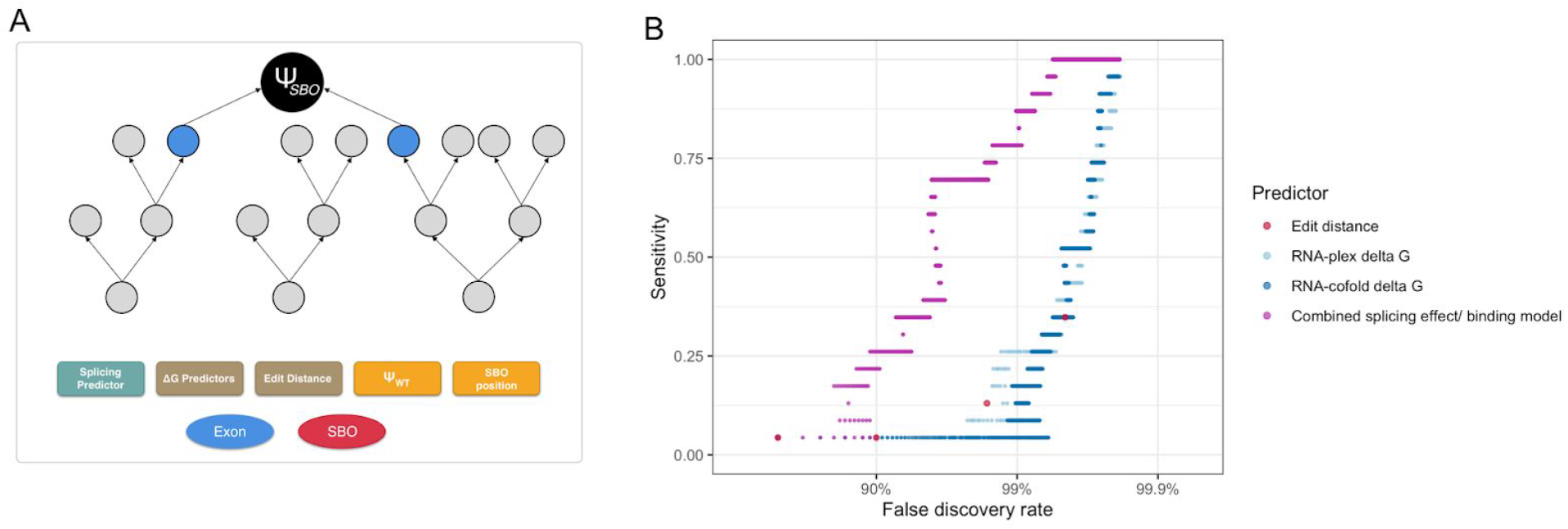
(A) Illustration of the gradient-boosted tree model trained to prioritize *in silico* hits at an edit distance of five or lower. (B) Performance when evaluated on a test set of unseen SBOs and exons. Significant splicing changes with dPSI greater than 0.2 were labelled as positives.

## Discussion

Steric-blocking oligos designed to hybridize to one site in the transcriptome can cause off-target effects by binding to near-complementary sites. Our results suggest that expression changes are more common than splicing changes, but also that only off-target splicing effects are predominantly hybridization-dependent and more reproducible. Changes in transcript levels may be driven by other off-target mechanisms, or indirect transcriptional effects of on-target effects, or confounders such as batch effects. Technical factors such as transfection efficiency could also affect the reproducibility of differential expression events. Given the enrichment of splicing effects at low edit distances to in-frame exons, we would expect that there is a set of genes downregulated due to off-target binding to an out-of-frame exon, but this effect is not clear from the data due to a high background rate of differential expression events.

Limited reproducibility and the lack of a clear, hybridization-driven mechanism suggests that differential expression events need to be replicated before they can be considered true off-target effects. Differential splicing results, especially when of large effect, are more reproducible, although replication experiments could still be necessary to accurately detect smaller splicing changes.

Unlike for gapmers, hybridization to an off-target region is not enough for an SBO to cause an effect (Obad et al., 2011). Intronic binding sites rarely lead to a change in splicing, and even the majority of exonic binding sites do not cause a large effect, making it difficult to predict off-target events based on binding affinity alone. A previous study of two gapmers found 54% of gene expression changes to occur at an edit distance of zero or one, with a false discovery rate of 72% (Yoshida et al., 2019). By contrast, we found an edit distance cutoff of one to the exon body to result in a sensitivity of 3.4% at a false discovery rate of 91% for differential splicing events. Differential expression events are even more difficult to predict from *in silico* screens, as there is no clear association with edit distance.

These results suggest that *in silico* methods that consider all binding sites below a certain edit distance as potential off-target hits are of limited use for SBOs, as they are likely to only capture a small fraction of all off-target events and result in a high false discovery rate. Empirically searching for these events using RNA-seq may be a superior approach to detecting these events, although follow-up experiments may be needed to replicate the findings. Better predictors that incorporate both binding affinity and splicing effects could also improve the utility of *in silico* methods.

Our study is limited to PS-MOE SBOs in three different cell types, and does not investigate the effect of different chemistries or delivery methods on off-target hybridization. Larger studies that include *in vivo* experiments will be needed to understand the extent of hybridization-dependent off-target effects for therapeutic compounds.

## Acknowledgements

We would like to thank the staff of The Centre for Applied Genomics for library preparation and sequencing.

## Supplementary Figures

**Supplementary figure 1:**
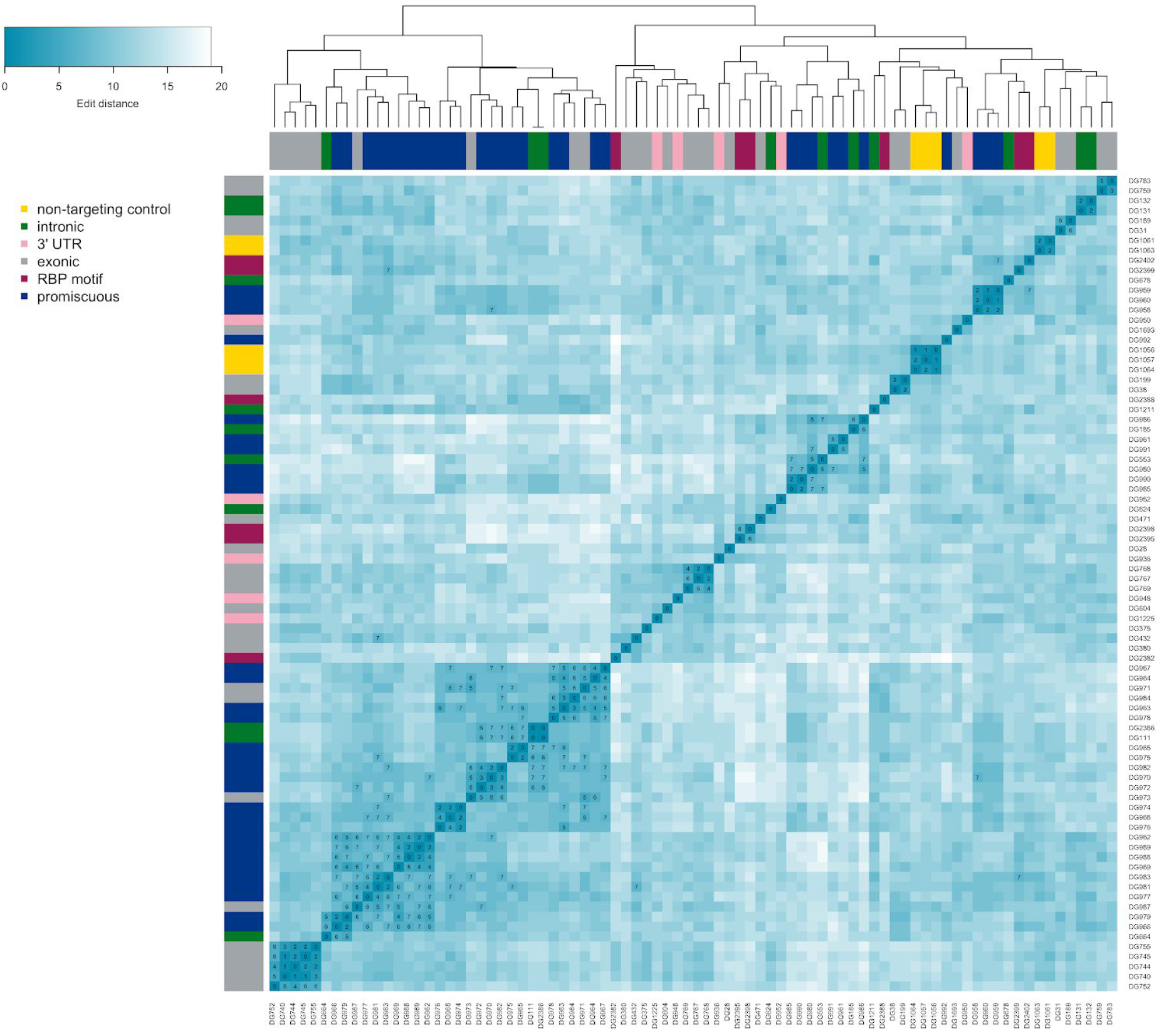
Pairwise edit distances between all SBOs included in the study. The edit distance is indicated by the colour and additionally shown as a text label for edit distances less than or equal to seven. DG2386 is a biotin-labelled version of DG111.

**Supplementary figure 2:**
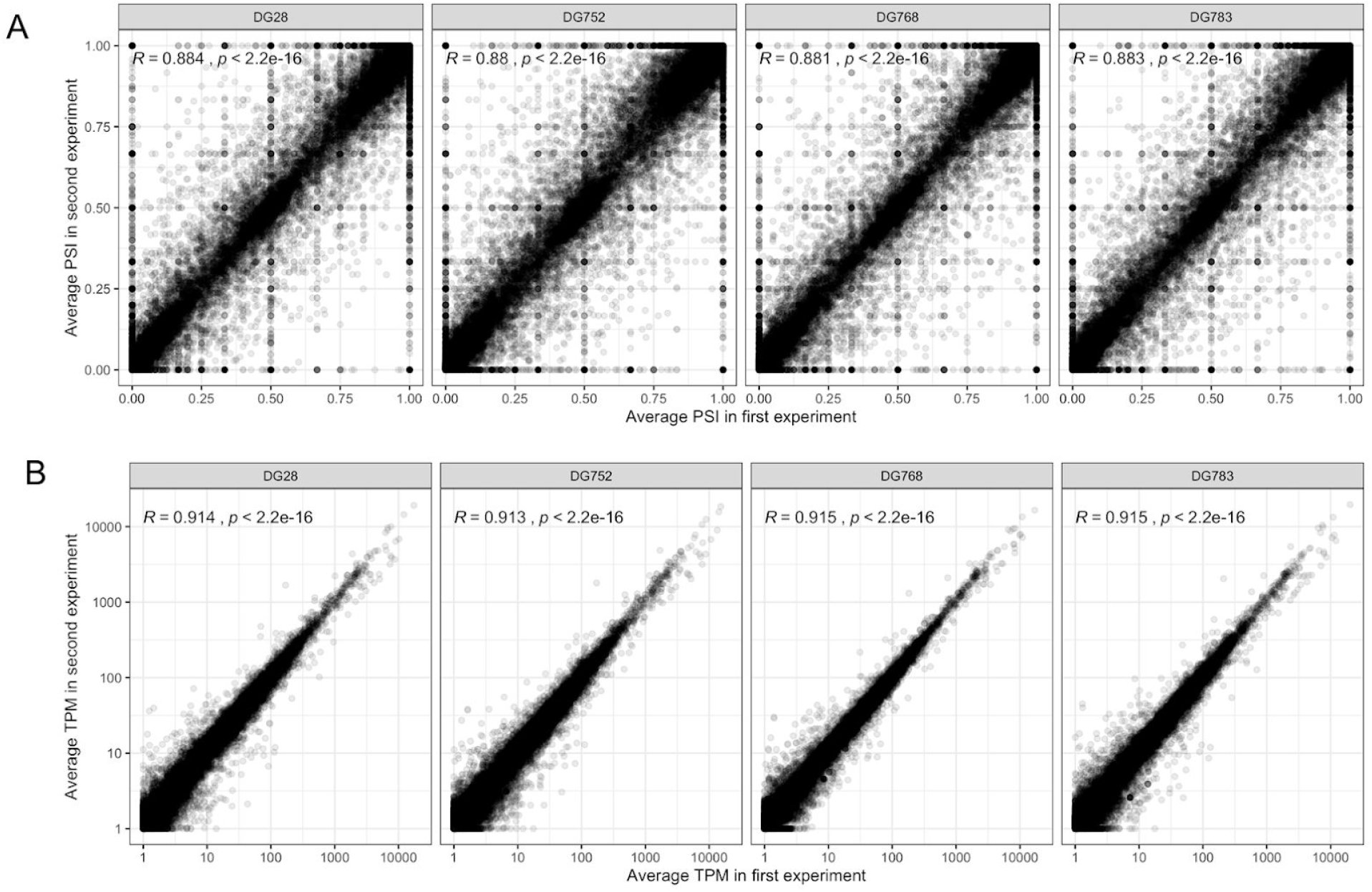
Reproducibility of exon PSI-estimates (A) and transcripts per million (B) between repeated experiments. Spearman correlation is shown for each SBO.

**Supplementary figure 3:**
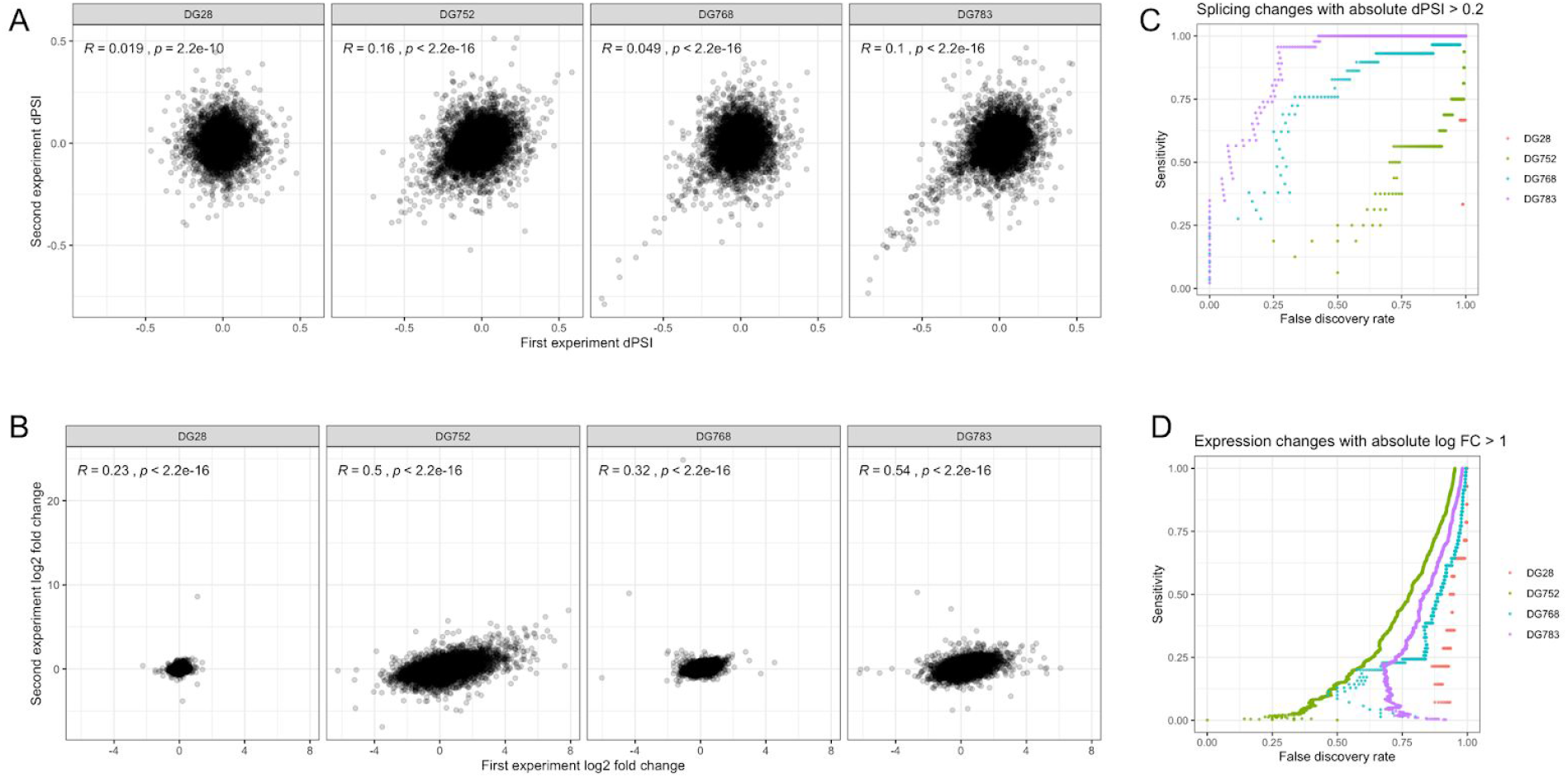
Reproducibility of differential splicing and expression estimates in repeat experiments. (A) Spearman correlation between dPSI estimates. (B) Spearman correlation between log fold changes. (C) Performance when using observed absolute dPSI in the first experiment to predict significant changes with absolute dPSI > 0.2 in the replication experiment. (D) Performance when using observed absolute log fold change in the first experiment to predict significant changes with absolute log fold change > 1 in the replication experiment.

**Supplementary Figure 4:**
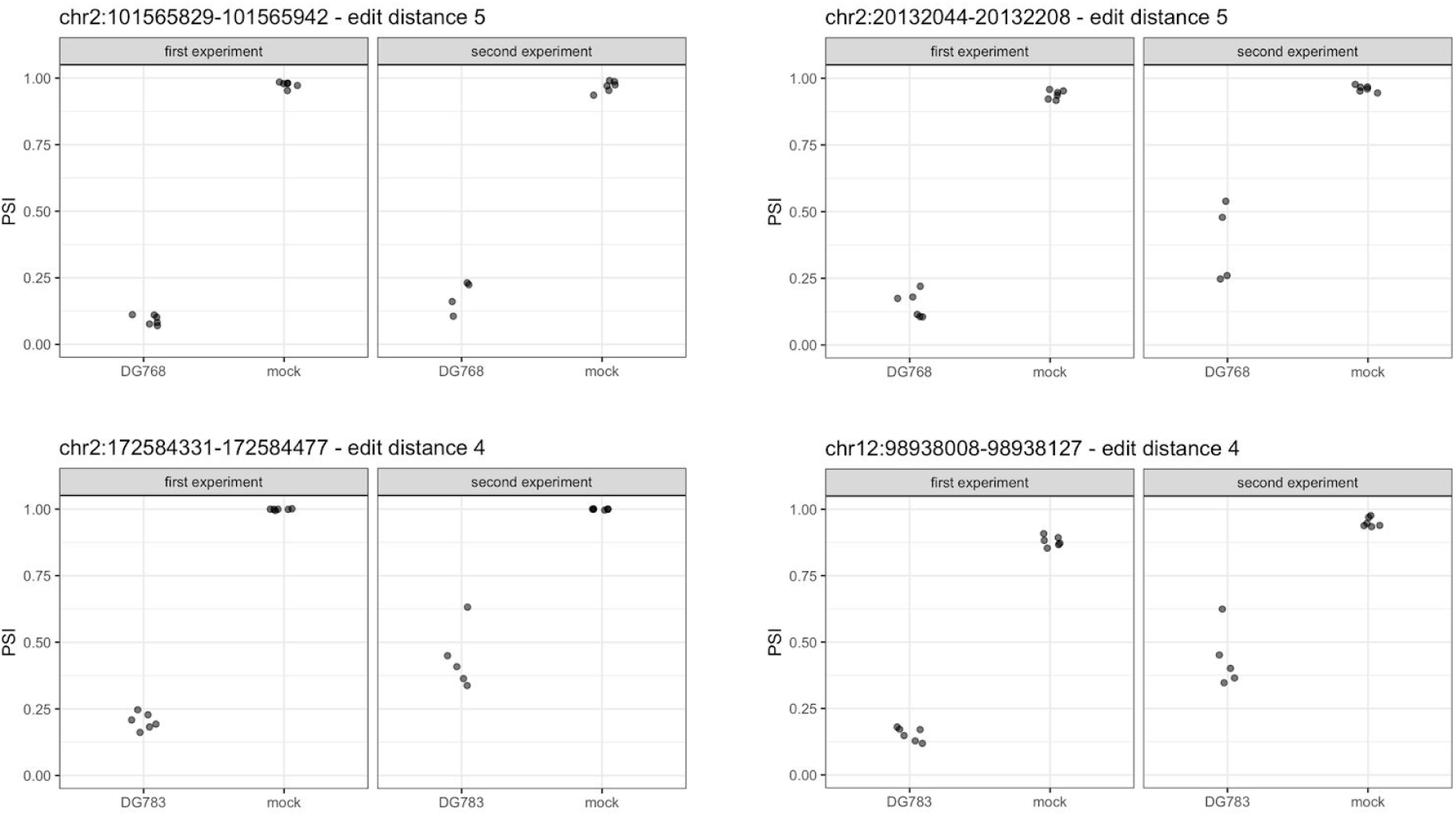
Examples of replicated differential splicing events at high edit distances for two SBOs.

**Supplementary figure 5:**
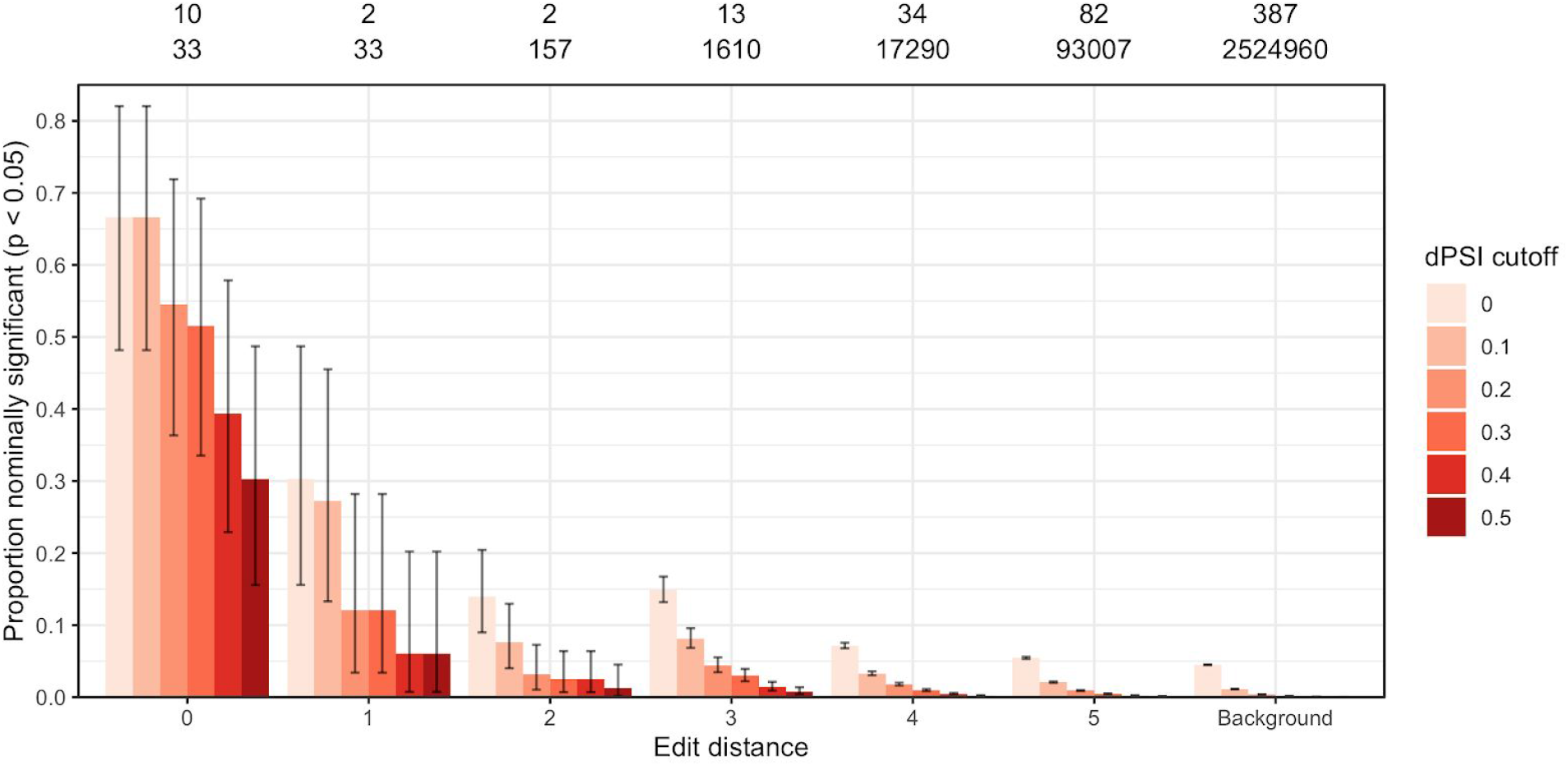
Enrichment of differential splicing events by edit distance for the subset of 46 SBOs used for differential expression.

**Supplementary figure 6:**
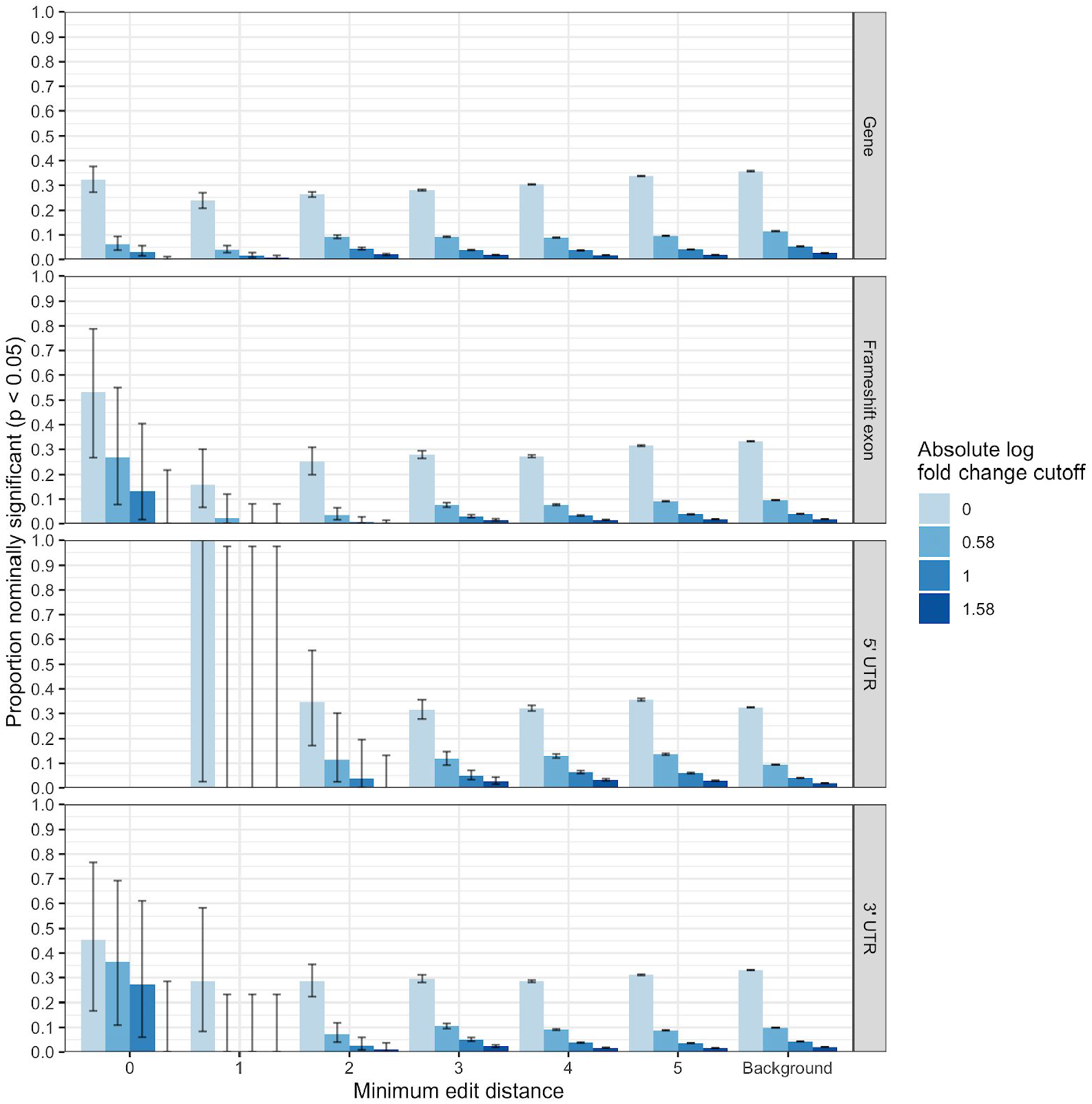
Gene expression changes as a function of edit distance, for different categories of hits. Both upregulation and downregulation events were included.

**Supplementary figure 7:**
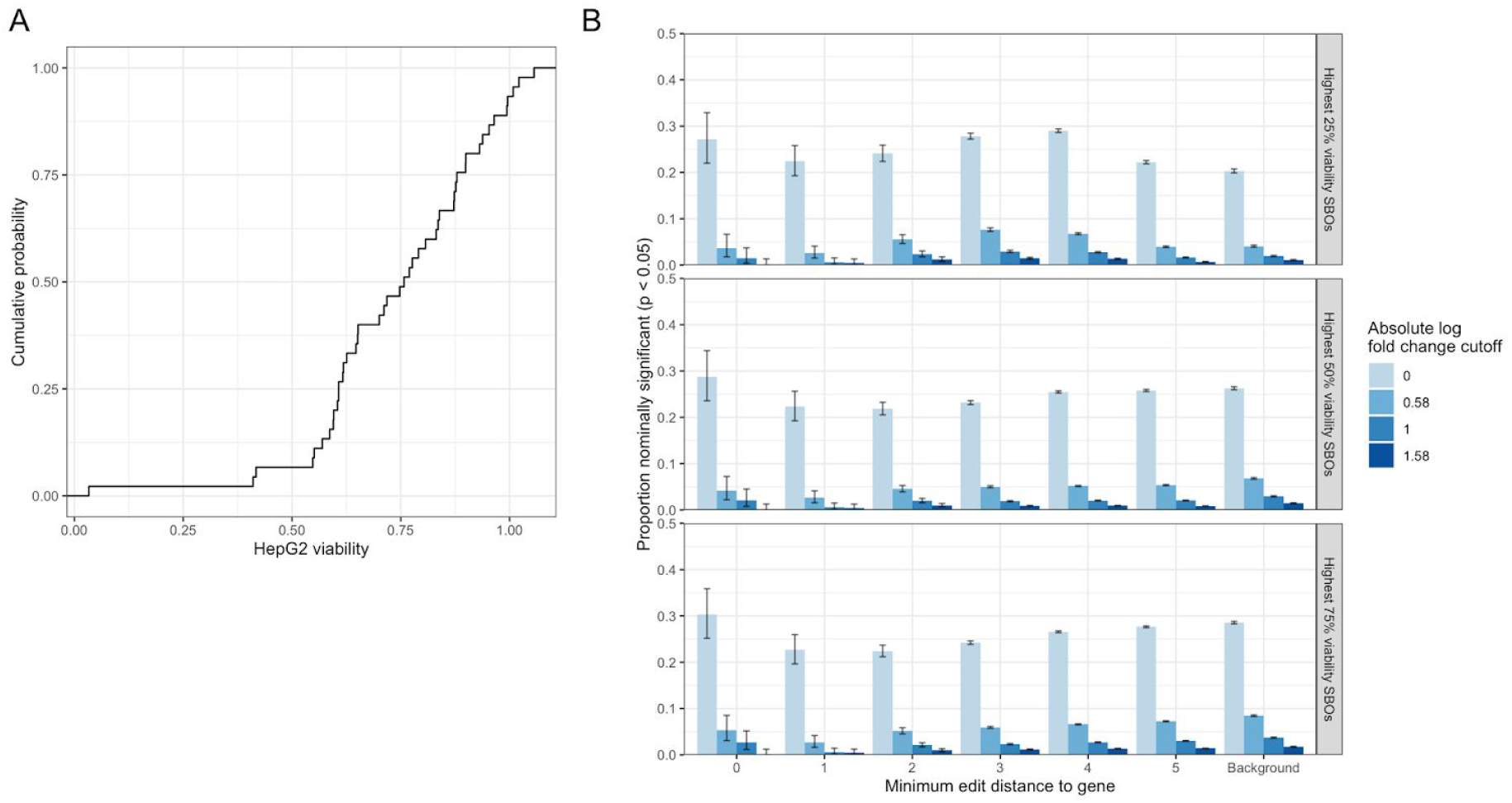
(A) Distribution of normalized cell line viability scores for SBOs included in differential expression analysis. (B) Differential expression events by edit distance to full gene, broken down by absolute log fold change, for highest 75%, 50%, and 25% viability SBOs.

**Supplementary figure 8:**
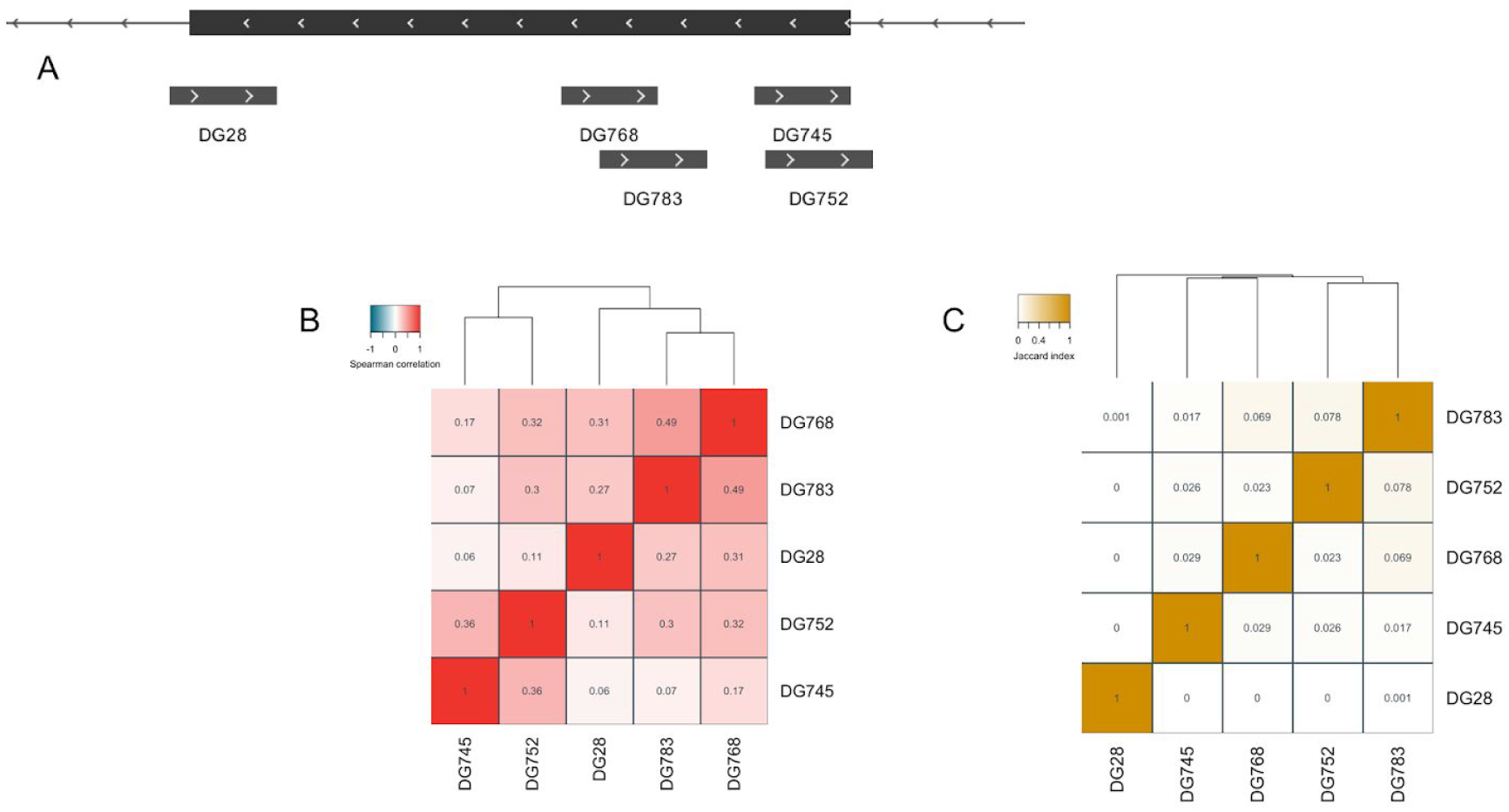
(A) Hybridization intervals of five SBOs targeting the same exon. (B) Spearman correlation of log fold changes between all pairs of SBOs. (B) Overlap of significant (q < 0.05) differential expression events with absolute log2 fold change greater than 1, for all pairs of SBOs.

**Supplementary figure 9:**
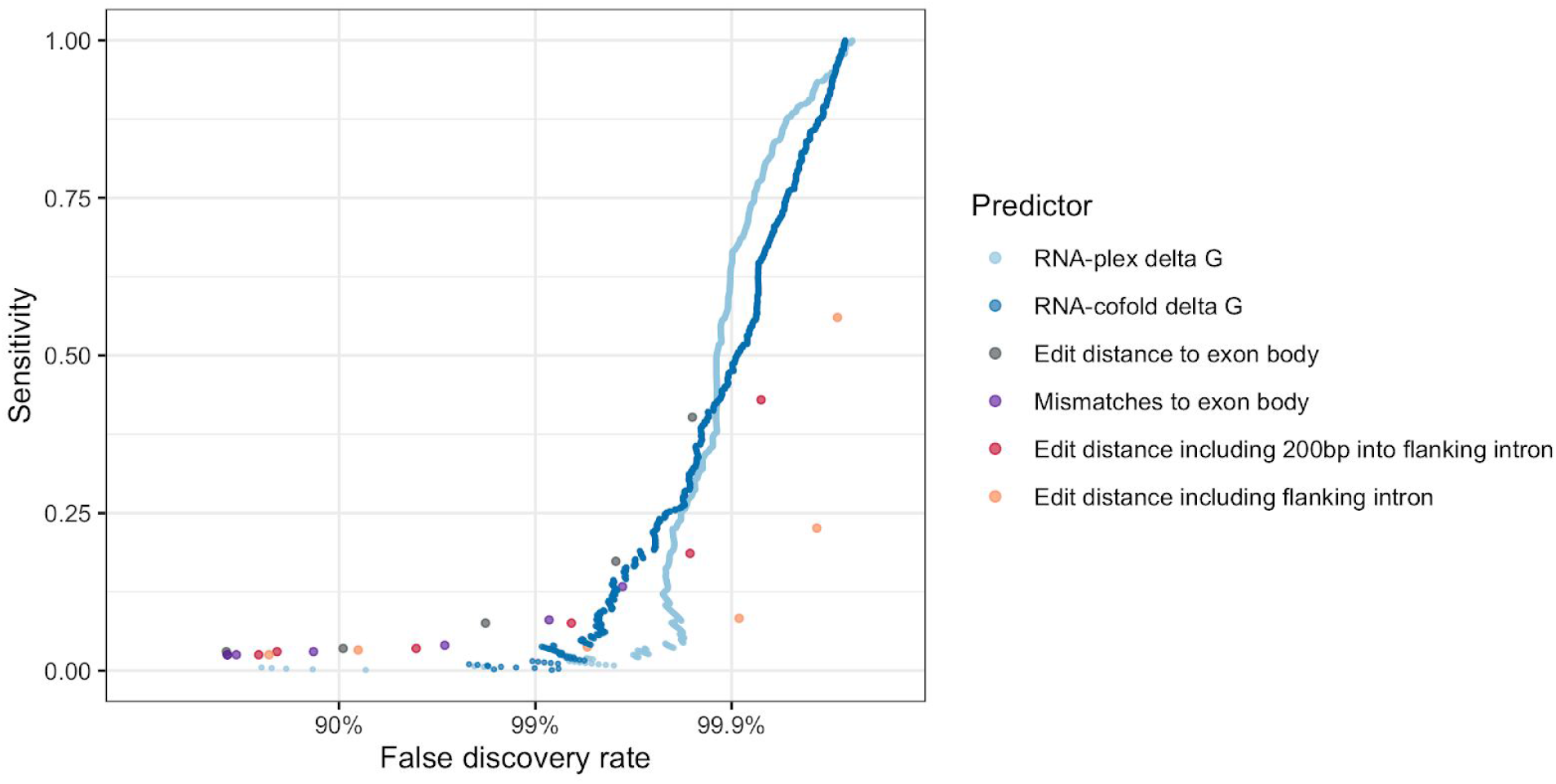
Performance of different predictors at identifying off-target splicing events with a change in dPSI of at least 0.5 when removing the 31 SBOs designed to have complementarity to several parts of the transcriptome.

## Supplementary Tables

**Supplementary table 1**

RNA-seq experiments included in the study.

**Supplementary table 2**

Off-target differential splicing events with q-value < 0.05 and absolute dPSI > 0.5.

**Supplementary table 3**

Off-target differential expression events with q-value < 0.05 and absolute log fold change > 1.

